# Onset, Time to Recurrence, and Recurrence Risk Factors of Myocardial Infarction and Ischemic Stroke: 10-Year Nationwide one-Million Population Database

**DOI:** 10.1101/253617

**Authors:** Kang Jae Jung, Hyun Sun Lim, Hyoung Seop Kim

## Abstract

**Background and Purpose:** We aimed to determine the differences in the pathophysiology of myocardial infarction (MI) and ischemic stroke (IS) and present their basic epidemiologic data for public health policy.

**Methods:** We included patients with a history of admission with diagnostic codes of MI and IS in the National Health Information Database (NHID) of the National Health Insurance Service (NHIS) Sample Cohort 2002-2013. We investigated the time to primary and secondary events, difference in incidence based on sex, time interval for the secondary event following the primary event in MI and IS, and the relative risk of recurrent events compared with that of primary events.

**Results:** The mean age of onset of IS was significantly higher than that of MI (65.67±12.25 vs. 60.4±13.07 years; *P*<0.05). The mean period from primary IS to secondary IS was significantly shorter than that from primary IS to secondary MI (2.09±2.80 vs. 5.35±2.90 years; *P*=0.016). The mean period from primary MI to secondary MI was significantly shorter than that from primary MI to secondary IS (2.30±2.75 vs. 5.16±3.26 years; *P*<0.001). The incidence of IS in men was significantly higher than that in women. The relative risk of recurrent IS and MI was greater than that of the primary event. In patients younger than 70 years, the incidence of MI was higher than that of IS, while in those above 70 years it was lower (*P*<0.05).

**Conclusions:** The age at onset, time to recurrence, sex, and relative risks are significantly different between ischemic stroke and myocardial infarction, suggesting that the underlying pathophysiologic mechanisms might be different between the two.

## Introduction

Despite the dramatic decline in the mortality of stroke and myocardial infarction (MI) owing to sustained therapeutic advances over the past several decades, MI and stroke have been the first and third most common cause of mortality in the western countries to date, respectively,^1^ and stroke is a more important cause of mortality and long-term disability.^2, 3^

It is generally accepted that atherosclerosis is a major cause of MI and ischemic stroke (IS).^4^ However, MI is mainly caused by the occlusion of a large coronary artery, whereas IS can occur owing to the occlusion of cerebral arteries due to atherosclerosis or embolus but shows many heterogeneous features.^5, 6^ Despite continuous improvements in therapeutic methods, MI and IS recurrence rates within 5 years have been reported at 21–33% and 25-37%, respectively.^7, 8^ Furthermore, the primary event acts as a risk factor and affects mutual recurrence.^9^

Therefore, for preventing primary and secondary overall cardio-cerebrovascular disease, it is necessary to investigate the epidemiologic features of, both, IS and MI and the factors associated with recurrence.

However, to date, there have been some reports of epidemiology, treatment, and prevention of either IS or MI, but few comprehensive studies on both diseases in a single cohort have been reported.

In this study, we included the patients with a history of admission with diagnostic codes of MI and IS in a 2.2% random sample cohort of the total Korean population from 2002 to 2013 (the National Health Information Database (NHID) of the National Health Insurance Service (NHIS) Sample Cohort; NHID-NHIS 2002-2013). We aimed to investigate the basic epidemiologic data for public health and clarify the differences in the pathophysiology of MI and IS.

## Methods

### Statement of Ethics

This study was reviewed and approved by the Institutional Review Board of our hospital adhered to the tenets of the Declaration of Helsinki. Informed consent was waived. All authors contributed to the study design, interpretation of the results, and made the decision to submit the manuscript for publication. No commercial support for this study was obtained.

### Database

This retrospective study used NHID-NHIS 2002-2013, which was released by the Korean National Health Insurance Service in 2015. The data comprises of 1,025,340 nationally representative random subjects, amounting to approximately 2.2% of the entire population (46,605,443) in 2002. To distinguish the patients from those with previous illnesses, we excluded 14,520 patients with MI and IS in 2002 and the remaining patients were followed up from January 2002 to December 2013 (Figure 1).

**Fig 1.**
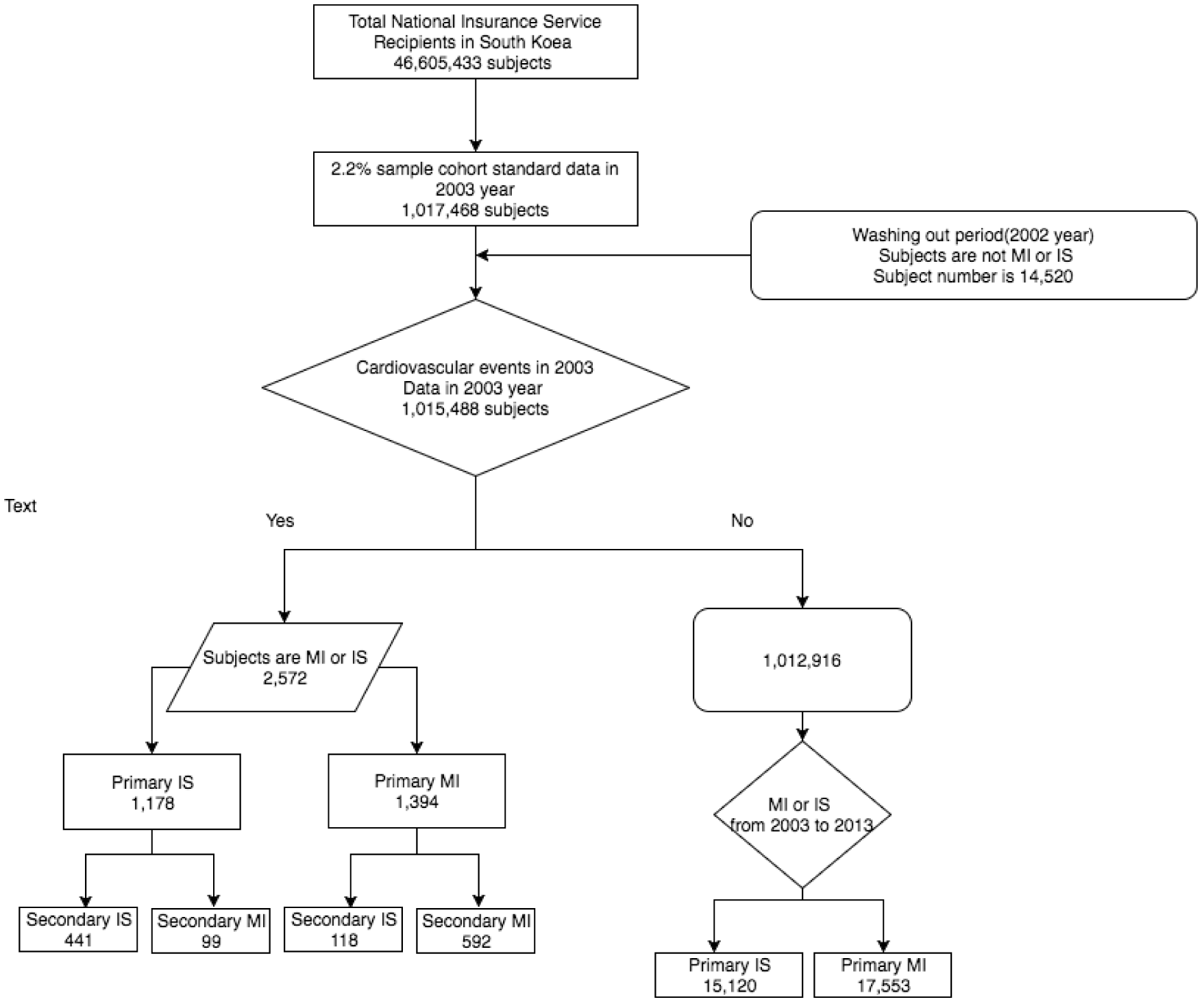
Flowchart for the Selection of Subjects

The data were generated by the NHIS using a systematic sampling method to generate a representative sample of all Korean residents. This database includes all medical claims filed from January 2002 to December 2013 (Figure 1). We extracted the subjects with cerebral infarction (I63) and myocardial infarction (I20-1) using the International Classification of Disease (ICD-10) in NHIS-NSC 2002-2013. Each primary group of MI and IS comprised subjects who were admitted to the hospitals due to the primary events of ischemic stroke and myocardial infarction, respectively, and then the subgroups were divided on the basis of the successive secondary event. Additionally, the subjects were classified into four subgroups: MIMI (secondary MI after primary MI), MIIS (secondary IS after primary MI), ISMI (secondary MI after primary IS), and ISIS (secondary IS after primary IS).

### Statistical analysis

Statistical analysis was performed using SAS system for Windows, version 9.4 (SAS Inc., Cary, NC, USA). Statistical significance was set at 0.05 or less. Independent t-test was performed to verify the differences between MI and IS according to the mean age of onset, the time interval between primary and secondary event, and sex. A chi-square test was used to inspect the relationship between 5-year interval and a primary event in the two groups (MI vs. IS). Age-adjusted incidence rates were calculated and expressed in terms of per 100,000 person-years by the direct method using the standard population derived from the population of Korea in each year. The incidence rate of the difference between IS and MI was investigated using logistic regression analysis.

To compare the incidence of secondary events according to the primary events, we classified and compared the ISIS and ISMI groups with the MIIS and MIMI groups, respectively, by estimating the relative risk of incidence (RR) with 95% confidence intervals (CI).To identify the effect of a primary event on the recurrence, we also obtained RR of the incidence of secondary events to that of the primary events without a secondary event after 2003.

## Results

The mean age of occurrence of IS was significantly higher than that of MI (65.67 ± 12.25 vs. 60.46 ± 13.07 years, mean ± SD, *P*< 0.01).

The mean period of ISIS was significantly shorter than that of ISMI (2.09 ± 2.80 vs. 5.35 ± 2.90 years, P = 0.016). The mean period of MIMI was significantly shorter than that of ISIS (2.30 ± 2.75 vs. 5.16 ± 3.26 years, P < 0.001). However, there was no significant difference between the mean period of MIIS and ISMI groups.

There was no significant difference in the incidence of IS between men and women but the incidence of MI was significantly higher in men than in women (Table 1).

**Table 1.** The Difference in Primary Events between the Sexes.

|  |  | Male | Female | P |
| --- | --- | --- | --- | --- |
| Total Population number |  | 508,178 | 507,310 | 1,015,488 |
| IS | Subjects (n) | 600 | 578 | 0.5405 |
|  | % | 0.06 | 0.06 |  |
| MI | Subjects (n) | 781 | 613 | <.0001 |
|  | % | 0.08 | 0.06 |  |

$\chi^2$ analysis was performed to compare the sexes

The incidence of ISIS was higher than that of ISMI (RR, 4.45; CI, 3.64-5.45) and the incidence of MIMI was higher than that of MIIS (RR 5.01; CI, 4.18-6.03) (Table 2).

**Table 2.**
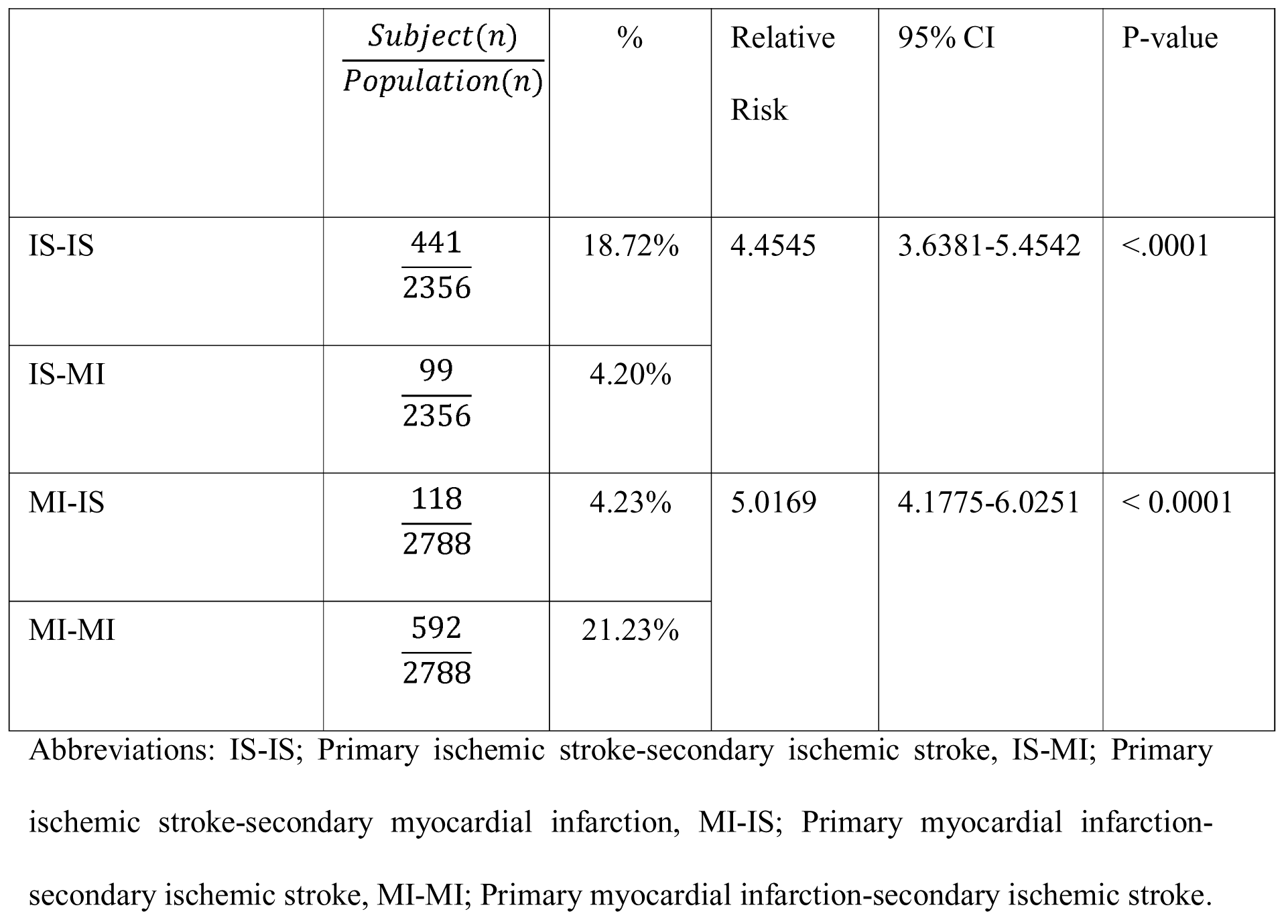
The Relative Risk of a Secondary Event after a Primary Event.

The risk of recurrent IS in patients with a previous primary cardio-cerebrovascular event in 2003 was higher than that in patients with primary IS after 2004 (RR 14.56; CI 13.51-15.69). In addition, the risk of recurrent MI in patients with a previous primary cardio-cerebrovascular event in 2003 was higher than that in patients with primary MI after 2004 (RR 15.50; CI, 14.52-16.55) (Table 3).

**Table 3.** The Relative Risk of Secondary Events of MI and IS after a Primary Event and the Incidence of a Primary Event.

| | $\frac{Subject(n)}{Population(n)}$ | % | Relative Risk | 95% CI | P-value |
| --- | --- | --- | --- | --- | --- |
| Secondary IS after primary cardiovascular Event | $\frac{559}{2572}$ | 21.73% | 14.5600 | 13.5076-15.6944 | <.0001 |
| Primary IS Event (2003-2013) | $\frac{15120}{1012916}$ | 1.49% | | | |
| Secondary MI after primary cardiovascular Even | $\frac{691}{2572}$ | 26.87% | 15.5035 | 14.5216-16.5518 | < 0.0001 |
| Primary MI Event (2003~2013) | $\frac{17553}{1012916}$ | 1.73% | | | |

A significant statistical difference was found between the incidences of, both, primary IS and MI with age (P <0.001). On logistic regression analysis, the interaction effect between age and the disease (IS and MI) was significant (P < 0.001) and the incidence of MI was higher than that of IS in patients younger than 70 years while the incidence of IS was higher than that of MI in patients aged above 70 years (Table 4) (Figure 2).

**Table 4.** The Incidences of both Primary cardiovascular events related with a 5-year interval

| Age (years) |  | <40 | 41-45 | 46-50 | 51-55 | 56-60 | 61-65 | 66-70 | 71-75 | 76-80 | >81 | Total |
| --- | --- | --- | --- | --- | --- | --- | --- | --- | --- | --- | --- | --- |
| IS* | N | 32 | 34 | 61 | 85 | 94 | 188 | 190 | 191 | 162 | 141 | 1,178 |
|  | % | 0.01 | 0.04 | 0.08 | 0.16 | 0.21 | 0.43 | 0.56 | 0.84 | 1.13 | 0.98 | 0.12 |
|  | N/100,000 | 5.18 | 36.36 | 78.96 | 157.05 | 211.33 | 434.60 | 560.64 | 838.90 | 1129.16 | 976.86 | 116.00 |
| MI* | N | 80 | 85 | 123 | 142 | 158 | 239 | 213 | 168 | 100 | 86 | 1,394 |
|  | % | 0.01 | 0.09 | 0.16 | 0.26 | 0.36 | 0.55 | 0.63 | 0.74 | 0.70 | 0.60 | 0.14 |
|  | N/100,000 | 12.96 | 90.90 | 159.21 | 262.37 | 355.20 | 552.50 | 628.50 | 737.87 | 697.01 | 595.81 | 137.27 |
| Total<br>Population | Number | 617,421 | 93,514 | 77,253 | 54,122 | 44,481 | 43,258 | 33,890 | 22,768 | 14,347 | 14,434 | 1,015,488 |
|  | % | 60.80 | 9.21 | 7.61 | 5.33 | 4.38 | 4.26 | 3.34 | 2.24 | 1.41 | 1.42 | 100.00 |

Abbreviation: N, Number
\*; P<0.001, A logistic regression analysis were performed..

**Fig 2.**
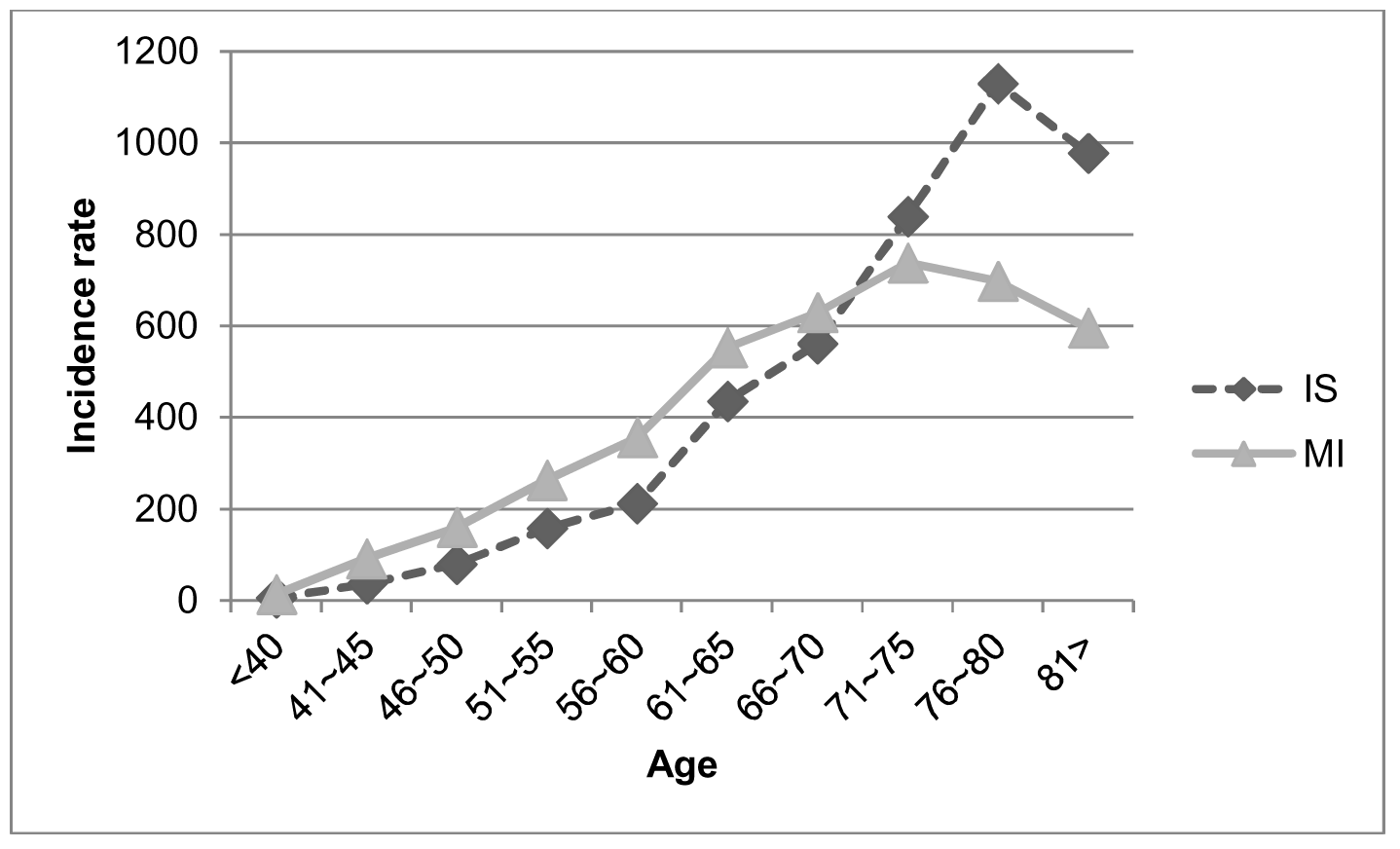
The Comparison between the incidence of Primary MI and IS (N/100,000) by 5-year intervals

## Discussion

The mean onset age of stroke and MI varies according to region and race, with that of stroke ranging from 64.1 to 71.5 years and MI ranging from 64.1 to 70.5 years.^10-19^

According to another Korea Stroke Registry from 2002 to 2010 (n=46,098), the mean onset age of stroke was 66.1±12.3 years and this is similar to the results of this study.^18^

In a single cohort study of 1,660 patients with IS and MI in the Dijon Vascular project, the mean onset age of IS was 7 years higher than that of MI (71.5±13.1 vs. 64.9±14.7) in men, but was similar in women.^10^ In our study, the mean onset age of IS was approximately 5 years higher than that of MI (65.67±12.25 vs. 60.46±13.07). The reason for this observation is unclear. However, it is believed that the pathophysiologic mechanisms of MI and IS are different. First, the major cause of MI is atherosclerotic changes in large vessels. If there is partial atherosclerotic occlusion of coronary arteries, MI can develop when there is an abrupt increase in the myocardial oxygen demand without total occlusion. IS can be also affected by the atherosclerotic changes in large vessels; however, if a total occlusion due to rupture of an atheromatous plaque does not occur, blood flow may be maintained and symptoms may not occur. IS in small vessels that supply subcortical areas, unlike in large vessels, can develop due to atheriolosclerotic pattern of necrosis or apoptosis of smooth muscle cells.^19^ Furthermore, there are many cases of asymptomatic IS in the subcortical areas of the brain. Second, in contrast to a coronary artery in MI, when a large cerebral artery gets occluded gradually, the development of multiple collateral vessels can maintain the cerebral circulation and it is considered to contribute to the delay in the onset of symptoms.^20^ Third, is already known, IS and MI have similar risk factors. However, hypercholesterolemia is more strongly associated with MI than with IS but hypertension is more strongly associated with IS than with MI. IS requires more time for the development of vessel damage and is considered to occur later in onset.^21^ Fourth, atrial fibrillation and the resulting embolic infarction also increases with age, which may be the reason for increasing IS with age.^22^ Finally, atrial fibrillation may occur after acute MI due to cardiac structural or electrical changes, which can result in secondary embolic infarction as well.^23^

According to another Korea Stroke Registry from 2002 to 2010 (n=46,098), the mean onset age of stroke was 66.1±12.3 years and this is similar to the results of this study.^18^

The incidences of IS and MI are generally known to be higher in men than in women.^18, 24-26^ In the OXVASC study,^20^ the incidences of stroke and MI in men through all ages were similar and the incidence of stroke was higher than that of MI in women aged above 75 years. In our study, the incidence of MI was higher in men than in women. Women have been reported to have a relatively lower blood pressure compared to men due to the influence of female sex hormones on the pressure-natriuresis relationship.^27^ However, blood pressure is more influenced by age rather than sex; therefore, gender (behavioral) factors are considered to be more important than sex (biologic) factors.^27^ In other words, although there are regional differences, in men it appears that there are more risk factors such as smoking, obesity, hypertension, and poor treatment participation for dyslipidemia.^26, 28^

Regarding the recurrence after a primary event, the relative risk of ISIS was higher than that of ISMI, and that of MIMI was higher than that of MIIS. In addition, the time interval from the primary event to the secondary event, was shorter in ISIS than ISMI, and that in MIMI was shorter than that in MIIS. A previous study reported that 62% of patients with primary IS showed coronary artery stenosis.^9^ In other words, if there is artherosclerosis of either a cerebral or coronary artery, there is a possibility that occlusion is already ongoing in other vessels and a secondary event is likely to be not only the recurrence of the original event itself (MI or IS) but also of another primary event. In previous studies, both MI and IS were found to have a high recurrence rate within 1 year, especially within 1 month. The development of a primary event after a different primary event (MIIS or ISMI) can be influenced by other factors such as systemic inflammation.^29, 30^ Although both diseases have similar risk factors, it is presumed that they may have different patterns of luminal stenosis of a vessel due to artherosclerosis and the collateral circulation accompanying it. Therefore, we can infer the reason that there is a high recurrence rate and a short recurrence period for the same secondary event as the primary event.

In a comprehensive analysis, the primary cardio-cerebrovascular event, including IS and MI, was associated with a higher incidence of secondary event than that of primary event.

Based on these results, to prevent a secondary event, there is need for overall consideration of the differences between MI and IS such as different developmental processes according to age, sex, and major risk factors. Furthermore, it is also believed that IS has a more heterogeneous pathogenic process such as cardiac embolism due to atrial fibrillation or a vascular malformation.

In our study, the age standardized incidence rate was higher in MI than in IS (116/100 000 in IS vs. 137/100,000 in MI). In most western countries, the incidence of MI was higher than that of IS.^31, 32^ Our study showed similar results as previous studies. Furthermore, it also showed that the incidence increased steeply with age and was highest in the 76–80-year age group for IS and 71–75-year age group for MI. The relative incidence of IS was significantly higher than that of MI in patients aged over 70 years but that of MI was higher than that of IS in those aged under 70 years. Similar to our study, the DIVA study showed that the incidence of MI was higher than that of stroke in patients aged less than 65 years but that of stroke was higher than that of MI in those aged over 65 years.^10^

## Study limitation

In the present study, IS and MI were classified on the basis of the diagnostic code registered in accordance with the KCD codes and the admission history rather than by the universal critical diagnostic criteria. We analyzed the data of a huge sample retrospectively; therefore, it is difficult to clarify causality besides the phenomenological characteristics, i.e., in case of ischemic stroke, the causes of stroke cannot be differentiated between cardiogenic embolic infarction, ischemic stroke after subarachnoid hemorrhage, and a congenital vascular anomaly such as Moyamoya disease. Underlying conditions such as family history, smoking, alcohol consumption, comorbidities, and medication history were not defined in this study. In addition, our study did not reflect the effect of prophylactic procedures or medications such as coronary vessel stent, carotid artery stent, and anticoagulation or antiplatelet medications after the primary events.

## Summary

The onset age, recurrence interval, sex, and relative risk are significantly different between IS and MI. There could be different pathophysiologic mechanisms between the two diseases. Therefore, the primary and secondary preventive strategies for both the diseases should be prepared individually by age, sex and the primary event.

